# High-density amplicon sequencing identifies community spread and ongoing evolution of SARS-CoV-2 in the Southern United States

**DOI:** 10.1101/2020.06.19.161141

**Authors:** Ryan P. McNamara, Carolina Caro-Vegas, Justin T. Landis, Razia Moorad, Linda J. Pluta, Anthony B. Eason, Cecilia Thompson, Aubrey Bailey, Femi Cleola S. Villamor, Philip T. Lange, Jason P. Wong, Tischan Seltzer, Jedediah Seltzer, Yijun Zhou, Wolfgang Vahrson, Angelica Juarez, James O. Meyo, Tiphaine Calabre, Grant Broussard, Ricardo Rivera-Soto, Danielle L. Chappell, Ralph S. Baric, Blossom Damania, Melissa B. Miller, Dirk P. Dittmer

## Abstract

SARS-CoV-2 is constantly evolving. Prior studies have focused on high case-density locations, such as the Northern and Western metropolitan areas in the U.S. This study demonstrates continued SARS-CoV-2 evolution in a suburban Southern U.S. region by high-density amplicon sequencing of symptomatic cases. 57% of strains carried the spike D614G variant. The presence of D614G was associated with a higher genome copy number and its prevalence expanded with time. Four strains carried a deletion in a predicted stem loop of the 3’ untranslated region. The data are consistent with community spread within the local population and the larger continental U.S. No strain had mutations in the target sites used in common diagnostic assays. The data instill confidence in the sensitivity of current tests and validate “testing by sequencing” as a new option to uncover cases, particularly those not conforming to the standard clinical presentation of COVID-19. This study contributes to the understanding of COVID-19 by providing an extensive set of genomes from a non-urban setting and further informs vaccine design by defining D614G as a dominant and emergent SARS-CoV-2 isolate in the U.S.

## Introduction

The current COVID-19 pandemic is an urgent public health emergency with over 112,000 deaths in the United States (U.S.) alone. COVID-19 is caused by infection with the severe acute respiratory syndrome coronavirus-2 (SARS-CoV-2). The typical symptoms for COVID-19 may include the following: fever, cough, shortness of breath, fatigue, myalgias, headache, sore throat, abdominal pain, and diarrhea (Wu et al., 2020; Zhou, F. et al., 2020; Zhu et al., 2020). Patients admitted to the hospital generally have pneumonia and abnormal chest imaging (Bhatraju et al., 2020; Chen et al., 2020). COVID-19 is also associated with other complications, including acute respiratory failure and acute respiratory distress syndrome, which appear to be a significant predictor of mortality. Severe COVID-19 is disproportionately observed in the elderly and individuals with underlying comorbidities. COVID-19 has not similarly impacted children (Guan et al., 2020; Team, 2020; Verdoni et al., 2020; Xu, Y. et al., 2020), a rather atypical pattern for many viral respiratory diseases; however, other SARS-CoV-2 disease manifestations, such as Kawasaki disease, are emerging in this group.

The first reported SARS-CoV-2 clusters appeared in the Wuhan province in China and have since rapidly spread across the world (Li et al., 2020; Wu et al., 2020; Zhu et al., 2020). The primary means of transmission is by oral secretions, though viral RNA has also been detected in blood, stool, and semen (Kim et al., 2020; Zou et al., 2020). Social distancing, rapid case ascertainment, physical barriers, and quarantine of individual infected persons have proven successful in limiting the impact of COVID-19. For these public health measures to remain effective and sustainable, it is important to understand the pathways of transmission through aggressive contact tracing and virus testing. Of high concern with regards to SARS-Cov-2 is that the virus may be shed prior to the onset of clinical symptoms, at late times after the cessation of clinical symptoms, and by asymptomatically infected persons (Arons et al., 2020; He et al., 2020; Hijnen et al., 2020; van Doremalen et al., 2020; Wolfel et al., 2020; Xu, K. et al., 2020). While antibody testing identifies patients with prior exposure (Long et al., 2020), only targeted nucleic acid amplification testing (NAT) or SARS-CoV-2 antigen detection can identify actively transmitting individuals.

The SARS-CoV-2 genome shares 79.6% sequence identity with SARS-CoV, the causative agent of SARS in 2002. It shares 96% sequence identity with a bat coronavirus BatCoV, RaTG13 (Ceraolo and Giorgi, 2020; Lu, R. et al., 2020; Zhou, P. et al., 2020). SARS-CoV entry is determined by the spike protein ORF S (Wan, Y. et al., 2020). ORF S has many interaction surfaces and is the target of neutralizing antibodies. The S protein uses the human ACE2 (hACE2) as a receptor and is proteolytically activated by human proteases (Hoffmann et al., 2020; Shang et al., 2020). Comparative analysis shows that between SARS-CoV-2 and either SARS-CoV or bat-derived SARS-like coronavirus (bat SARS-CoV) (Andersen et al., 2020; Wu et al., 2020), the sequence identities are the least alike for spike protein gene (S). SARS-CoV-2 has a longer spike protein as compared to bat SARS-CoV, human SARS-CoV and middle east respiratory syndrome (MERS)-CoV (Lu, Roujian et al., 2020). Although SARS-CoV-2 only shares 79% identity with SARS-CoV at the whole genome scale, their spike protein receptor binding site sequences are more similar compared to bat SARS-CoV and MERS-CoV (Lu, Roujian et al., 2020). Residues at the receptor-binding site have evolved for better association with ACE2 compared to SARS-CoV (Wan, Yushun et al., 2020; Wrapp et al., 2020) and can be attributed to these molecular features: five of the residues critical for binding to ACE2 are different in SARS-CoV-2 as compared to SARS-CoV (Wan, Yushun et al., 2020; Wrapp et al., 2020) and a functional polybasic cleavage site (RRAR) is present at the S1/S2 boundary of the SARS-CoV-2 spike protein (Andersen et al., 2020; Walls et al., 2020). The polybasic cleavage site allows for effective cleavage by furin and other proteases which is important for viral infectivity (Letko et al., 2020). The additional proline may also result in O-linked glycans to S673, T678, and S686 that can be important in shielding key epitopes or residues (Andersen et al., 2020). Ascertaining if these key residues remain invariable as the pandemic progresses or if they evolve over time is crucial to ensure testing accuracy and rational vaccine design.

Initial phylogenetic analysis of human SARS-CoV-2 genomes established three major variant types worldwide (Forster et al., 2020). Clade B was derived from A by a synonymous T8782C mutation in ORF1ab; and a nonsynonymous C28144T mutation that changes a leucine to serine in ORF8 (Ceraolo and Giorgi, 2020; Forster et al., 2020). Clade C was derived from B by a nonsynonymous G26144T mutation that changes a glycine to valine in ORF3a. A and C types are mainly found in Europe and the U.S. B type is mainly found in East Asia. Other analyses arrived at different clades and unfortunately different naming conventions (Zhang et al., 2020). Additional clades have since been recognized, including clade G, which is defined by a non-synonymous single nucleotide variant (SNV) in spike protein at amino acid position 614. The most recent phylogenies are available from GISAID (GISAID, 2020; Shu and McCauley, 2017) and Nextstrain (Hadfield et al., 2018).

To provide finer granularity about biological changes during SARS-CoV-2 transmission, we employed next generation sequencing (NGS) as an independent screening modality. This allowed us to reconstruct the mutational landscape of cases seen at a tertiary clinical care center in the southeastern U.S. from the start of the U.S. epidemic on March 3, 2020, until past the peak of the first major wave of infections. The first case in North Carolina (NC) was reported on March 26, 2020. The samples cover the period when community spread in NC was established, and when the state-wide stay at home order was issued (March 30 – May 8, 2020).

SARS-CoV-2 testing remains limited in many countries due to a shortage of personal protective equipment, testing kits, and diagnostic capacity. The Centers for Disease Control (CDC) guidelines during the time of sampling prioritized patients with specific clinical symptoms (fever, cough, and shortness of breath) and curtailed testing to only a subset of all probable cases. Individuals not fitting the clinical criteria for testing, as well as asymptomatic individuals, were excluded. To evaluate if any cases were missed because of this triage algorithm, nasopharyngeal (NP) swabs for three groups of patients were evaluated: n=175 known SARS-CoV-2 positive NP samples, n=41 known SARS-CoV-2 NP negative samples, n=12 NP samples of unknown status, i.e. the patient had symptoms justifying sample collection but was not prioritized for clinical SARS-CoV-2 testing. “Testing by sequencing” was negative for all negative samples, less sensitive for weakly positive samples and uncovered 5 new cases among previously not tested cases. The index case in NC was linked to the U.S. outbreak in the state of Washington. Phylogenetic analyses established the dominance of the S protein D614G SNV among this population, which has been increasing over time through community spread and was introduced initially by a person returning from Europe.

## Methods

### Samples

This study used remnant samples of universal transport media (UTM) from provider-collected deep nasopharyngeal (NP) swabs (https://www.cdc.gov/coronavirus/2019-ncov/lab/guidelines-clinical-specimens.html) after their clinical purpose had been completed. The SARS-CoV-2 status of each sample was determined by a clinical NAT approved under EUA at The University of North Carolina at Chapel Hill Medical Center (UNCMC) McLendon Clinical laboratories. None of the samples carried any identifiers other than the date of testing. Hence, this research was considered part of the QA/QC effort to support clinical testing and classified as non-human subject research by local IRB.

### RNA isolation

250 *μ*L of virus transport medium (VTM) from flocked NP swabs were adjusted to 1.0 mL with 0.1% Triton X-100 (proteomics grade, VWR: 97063-864) diluted in 1X phosphate-buffered saline (PBS, Life Technologies, Catalog # 14190-144). Samples were incubated at room temperature for 30 minutes in a 2.0 mL screw cap tube (Genesee # 21-265), vortexing every 5 minutes for 15-second pulses. 200 *μ*L of the permeabilized sample was then processed using the Machery-Nagel NucleoSpin Virus kit (Macherey-Nagel, Catalog # 740983.50 and 740983.250).

Carrier RNA (poly-A salt) was added to the mixture to a final concentration of 9 ng/*μ*L. DNase digestion was performed post-column binding (Macherey-Nagel, Catalog #740963) at room temperature for 5 minutes at a final concentration of 40 ng/*μ*L. RNA was eluted from the column using 60 *μ*L of RNase-free water pre-heated to 70 °C. For each processing batch, a negative reagent control and a negative cell pellet control was used. The reagent control consisted of 250 *μ*L of 1X PBS instead of 250 *μ*L of the sample in UTM. The cell pellet controls used were stored at −80 &C before the emergence of SARS-CoV-2. The total number of cells used per negative control was 10^6^ and were treated identically and concurrently to the UTM and reagent control processed samples. The 60 *μ*L of eluted RNA was processed for sequencing and viral load as described below.

### Real-time qPCR

Relative viral genome copy number was ascertained by real-time qPCR using primers and procedures established by the CDC (Lu, X. et al., 2020). 30 *μ*L input RNA was subjected to hexamer-primed reverse transcription. 9 *μ*l cDNA was used for qPCR containing 125 nM for each primer and SYBR green as the method of detection on a Roche LC480II Lightcycler and crossing point (CP) values determined by automated threshold method.

### Sequencing

As a positive control, we used Genomic RNA from SARS-Related Coronavirus 2, Isolate USA-WA1/2020, as provided by BEI/ATCC. This reagent was deposited by the Centers for Disease Control and Prevention and obtained through BEI Resources, NIAID, NIH: NR-52285. All samples were sequenced using random hexamer/dT priming as provided by the Thermo SARS-CoV-2 AmpliSeq kit according to manufacturer’s recommendations on an IonTorrent Chef and IonTorrent S5 sequencer. The amplicons are tightly tiled and overlapping. Amplicon sizes ranged between 68 and 232 nucleotides after trimming of low-quality sequences and all primer sequences (125-275 before trimming).

### Bioinformatic analysis

Following primer trimming according to the manufacturer’s recommendations, additional, custom steps were added. Specifically, all sequences were quality trimmed using the bbduk script (arguments: qtrim=rl trimq=20 maq=20 minlen=40 tpe tbo) from bbmap version 37.36. Each trimmed sequence was analyzed using CLC Genomics Workbench version 11.0. The trimmed reads were mapped to the SARS-COV-2 reference sequence (Accession: NC_045512). From each mapping, the following was collected: a consensus sequence, sequence variants, and mapping coverage. The consensus sequence was extracted from the mapping by quality voting. Regions at or below a coverage threshold of 3 were considered low coverage and N’s were inserted for ambiguity. SNV were called using the CLC bio algorithm (Qiagen Inc.) for human genome SNV calling. The threshold for reporting was set at >90% frequency and a minimum coverage of 10-fold with balanced forward and reverse reads for all SNV. Targeted regions were determined via Thermo SARS-CoV-2 designed BED file and sequences with 1x coverage across more than 99% of the 237 SARS-COV-2 amplicons were considered complete sequences. Any sequences with 1x coverage between 5% and 99% were considered partial genomes. Partial genomes are included in the variant calling analysis. All consensus sequences derived from this study were manually curated to revert poly-nucleotide-tract mutations to the reference sequence.

### Phylogenetic reconstruction

Full-length, viral genome consensus sequences were aligned using MAFFT (Katoh and Standley, 2013) with a PAM200 / k =2 scoring matrix, gap open penalty of 1.53 and offset value of 0.123 as implemented in Genious (Genious Ltd) using n = 92 sequences. A neighbor-joining tree was constructed using Genious Tree Builder (Genious Ltd) with bat coronavirus strain TG13 (MN996532) as outgroup. The number of bootstrap replicates was 1,000 with a support threshold of > 50%. S protein sequences were analyzed using MEGA X version 10.1.7 (Kumar et al., 2018). Specifically, evolutionary history was inferred using the Neighbor-Joining method (Saitou and Nei, 1987). The optimal tree with the sum of branch length = 0.6557 is shown. The percentage of replicate trees in which the associated taxa clustered together in the bootstrap test (800 replicates) are shown next to the branches. The evolutionary distances were computed using the Maximum Composite Likelihood method (Tamura et al., 2004) and are in the units of the number of base substitutions per site. The rate variation among sites was modeled with a gamma distribution (shape parameter = 1). This analysis involved 96 nucleotide sequences. Codon positions included were 1st+2nd+3rd+Noncoding. All positions with less than 95% site coverage were eliminated, i.e., fewer than 5% alignment gaps, missing data, and ambiguous bases were allowed at any position (partial deletion option). There were 3,822 positions in the final dataset. Further sequences, including the NC index cases, as deposited by the State Health Department were provided by GISAID (GISAID, 2020).

### Statistical analysis

Further statistical analysis and visualization was conducted using R v 4.0.0. The code is available on bitbucket.

## Results

### Whole Genome SARS-CoV-2 sequencing through high-density Amplicons

UNCMC used one of two NATs to test for the presence of SARS-CoV-2 RNA, one laboratory developed test based on the protocol by Corman et al. (Corman et al., 2020) and the commercially available Abbott real-time SARS-CoV-2 assay, both under the EUA provision of the U.S. Food and Drug Administration. Both tests report the presence or absence of SARS-CoV-2 RNA. Remnant NP samples were subjected to targeted sequencing using the Thermo Fisher AmpliSeq SARS-CoV-2 assay and S5 Ion Torrent sequencing platform. Individual sequence reads were mapped to the SARS-CoV-2 reference sequence (NC_045512) and a strain-specific consensus sequence was generated and SNV recorded. The finished genomes are submitted to GenBank and GISAID and were named according to convention (Coronaviridae Study Group of the International Committee on Taxonomy of, 2020).

A total of n=175 known positive samples and positive control (full-length genomic RNA from strain SARS-CoV-2/human/USA-WA1/2020) were subjected to NGS. The number of mapped reads varied substantially across samples, reflecting the differences in the amount of virus per sample. The distribution of 10x coverage for all samples is presented in **Figure 1A.** As expected, more mapped reads yielded higher coverage. Of the 33 negative controls, none had >10^2^ total reads aligned. Of the positive samples, greater than 5*10^3^ total mapped reads were needed to obtain 1x coverage of the whole genome, a minimum of 3.1×10^4^ reads were needed to obtain >90% coverage at 10x. The number of reads aligned varied depending on the viral load, as determined by real-time qPCR using CDC primer N1, but not total RNA, as determined using RNAse P, of the samples (**Figure 1B)**. In this assay, any CP <35 for SARS-CoV-2 qPCR yielded reliable coverage, which increased linearly with viral load. At a CP ≥35 most positive samples still yielded reads that mapped to the target genome and thus allowed detection of SARS-CoV-2 sequences; however, the results were less consistent, and coverage was more variable. As expected, total RNA (measured by RNAse P) was not associated with sequencing coverage and varied considerably across samples, even though each sample used the same amount of virus transport medium (VTM).

**Figure 1:**
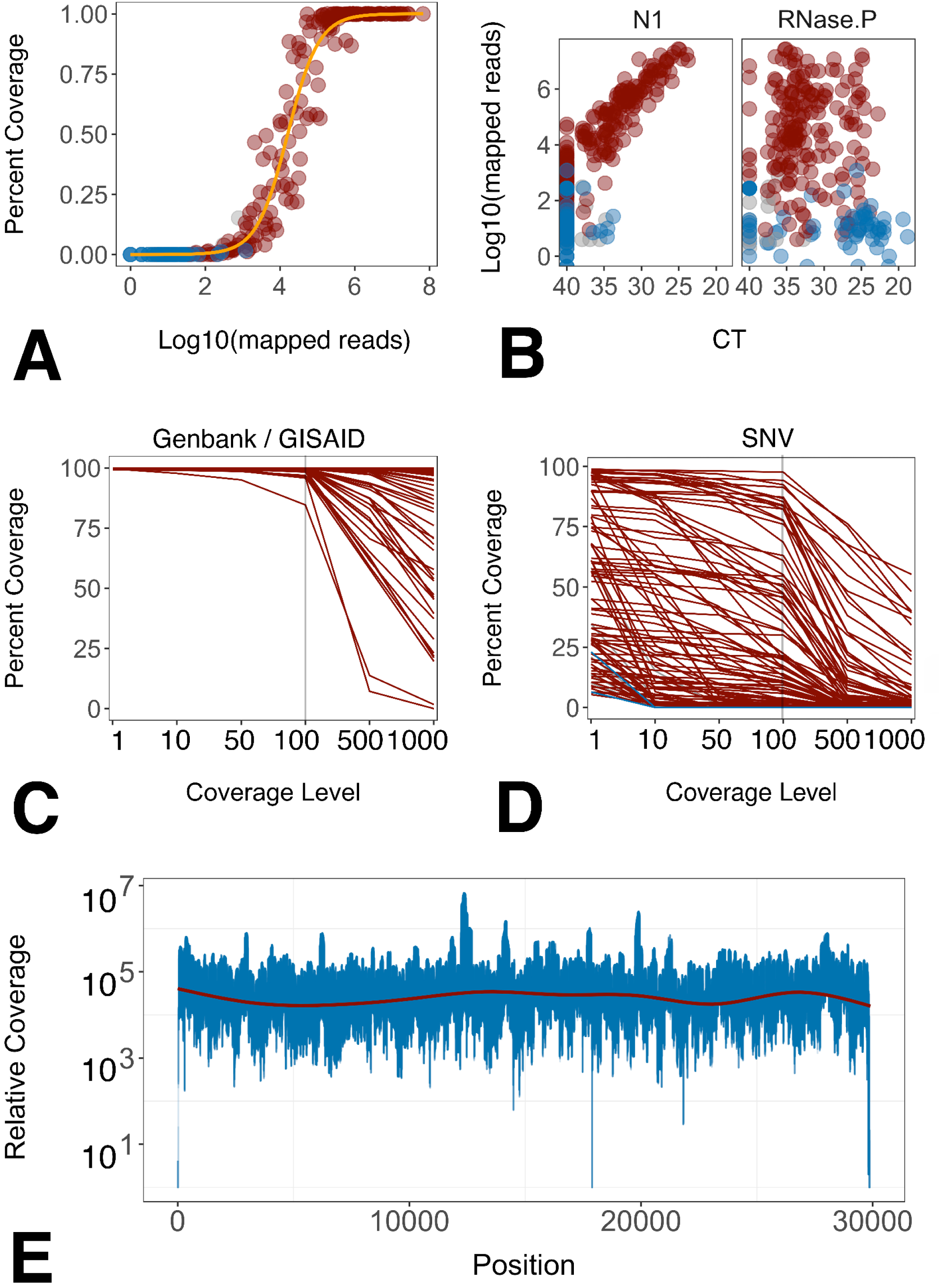
Analysis of sequencing coverage. A. Distribution of ≥10x coverage across all samples. Percent samples is shown on the vertical axis and log10 total mapped reads on the horizontal axis. Color indicate known positive (red), negative (blue), and unknown samples (gray). B. Relation between mapped reads, on the vertical axis, and relative viral load (N1) or total RNA levels on the horizontal axis. (C) Quality of samples submitted to GenBank GISAID. Percent genome covered on the vertical axis and coverage level at the horizontal axis. (D) Quality of additional samples used for SNV analysis. Percent genome covered on the vertical axis and coverage level at the horizontal axis. (E) Relative coverage of aligned reads per position of samples with median coverage >5000 reads is shown on the vertical axis for each position of SARS-CoV-2 as shown on the horizontal axis. Red line indicates a loess-fit of the data (n = 28 samples).

The coverage level distribution is shown in **Figure 1C and 1D**. Panel C represents all samples for which high-quality genomes were submitted to GenBank and GISAID. Panel D represents samples, with more variable complete coverage. These samples were nevertheless included in SNV calling, as the SNV algorithm relies on local coverage rather than overall coverage. As a result, the variant calls represent a conservative estimate of SNV distribution in this sample set. **Figure 1E** shows the per nucleotide coverage for all genomes with median coverage ≥5000x. Median coverage of >5000x was required to ensure > 99% genome coverage without a single amplicon dropout. The nucleotide composition of SARS-CoV-2 was largely balanced and did not contain repeats larger than sequencing length. Hence, coverage was continuous across the genome, except for the 5’ and 3’ untranslated regions (UTR). Targeted amplification using this primer set missed the first 42 nucleotides at the 5’ end and 29 nucleotides, starting at 29,843, at the 3’ end of the viral genome. These regions are conserved across most SARS-CoV-2 sequences in Genbank, many of which are themselves incomplete or known to suffer amplification bias (van Dorp et al., 2020). The limiting factor was not sequencing depth per se, rather, samples of low viral load failed in the targeted amplification step for individual amplicons. Samples with low viral load were re-sequenced.

Positive samples (n=33) were independently re-sequenced and yielded 251 high confidence SNV. No new SNV were uncovered upon resequencing; 180 SNV were confirmed and 71 SNV were lost upon pooling multiple sequencing runs for the same sample due the frequency dropping below 90%. Of the 71 SNV 50 possessed a majority vote matching the reference and 21 possessed a majority vote matching the prior SNV call. Target capture efficiency was verified using multiple dilutions and compared to unbiased RNAseq of the reference strain SARS-CoV-2/human/USA-WA1/2020 (**Supplemental Figure 1**). Targeted sequencing coverage was uniform over a 50-fold range of input RNA; it was higher than RNA seq, except in the terminal regions that were not covered by PCR amplicons. In some cases, as little as five microliters of VTM from a single swab had sufficient virus to obtain a full-length viral genome sequence at 1000x. This data is consistent with the astonishingly high reported genome copy numbers of SARS-CoV-2 in some cases (Yu et al., 2020) and demonstrates the principal suitability of “testing by sequencing” as a diagnostic option for SARS-CoV-2 and other rapidly evolving viruses.

Twelve samples were collected during the same calendar period from individuals presenting with respiratory complaints, but no indication for SARS-CoV-2 testing according to CDC guidelines. 5 of 12 yielded > 5% genome coverage (**Supplemental Figure 2**). The remainder had reads aligned only to regions of the genome that have low complexity; 2/12 had a sequence coverage distribution, at 57% and 34% respectively, consistent with the presence of the target virus. Three other samples had coverage of 20%, 13%, and 10%. At the time of study, SARS-CoV-2 testing guidelines were extremely restrictive due to a lack of supplies. Patients with clear clinical symptoms of COVID-19 were not tested but treated on the basis of clinical diagnosis alone, and patients with respiratory symptoms not exactly matching CDC/COVID-19 criteria were not tested either. None of the samples in this study originated from asymptomatic patients. Though the numbers of unknowns tested were small, the results suggest that limiting testing to narrowly defined case criteria misses a significant number of cases and thus transmission events.

### Sequence analysis reveals the presence of two clades of SARS-CoV-2

Putting individual sequences into context is key to understanding SARS-CoV-2 transmission. Sequencing identified n=139 samples with at least one high confidence SNV as compared to the reference sequence. Of these n=79 (57%) carried the S protein D614G SNV, a mutation implicated in higher pathogenicity of the virus (Becerra-Flores and Cardozo, 2020). Samples carrying the D614G SNV had higher SARS-CoV-2 genome loads as measured by CDC N3-primer directed real-time RT-qPCR for SARS-CoV-2 (p≤ 0.002 by Wilcox-Sign-Rank test). A similar, but not significant trend emerged using CDC N1-primer directed real-time RT-qPCR for SARS-CoV-2, but not for total RNA levels as measured by CDC RNAse P-directed real-time RT-qPCR (**Supplemental figure 3**). **Figure 2A** shows the SNV distribution of the data, color-coded by the week of collection. These data include high confidence SNV of genomes with < 99% coverage, whereas the phylogenetic reconstructions are only based on complete genomes (≥99% coverage) that were submitted to GenBank (and also present in GISAID). This SNV distribution was dominated by isolates representing clade A and some of clade B, the dominant clades in North America and Europe (Forster et al., 2020). The NC stay at home order was enacted on March 30, 2020, and the sample collection concluded on April 11, 2020, i.e. covering a period of unrestrained local spread. The SNV pattern is consistent with the idea that SARS-CoV-2 was introduced into NC by travelers from the continental U.S. and that this population was in equilibrium with the general population of the U.S.

**Figure 2:**
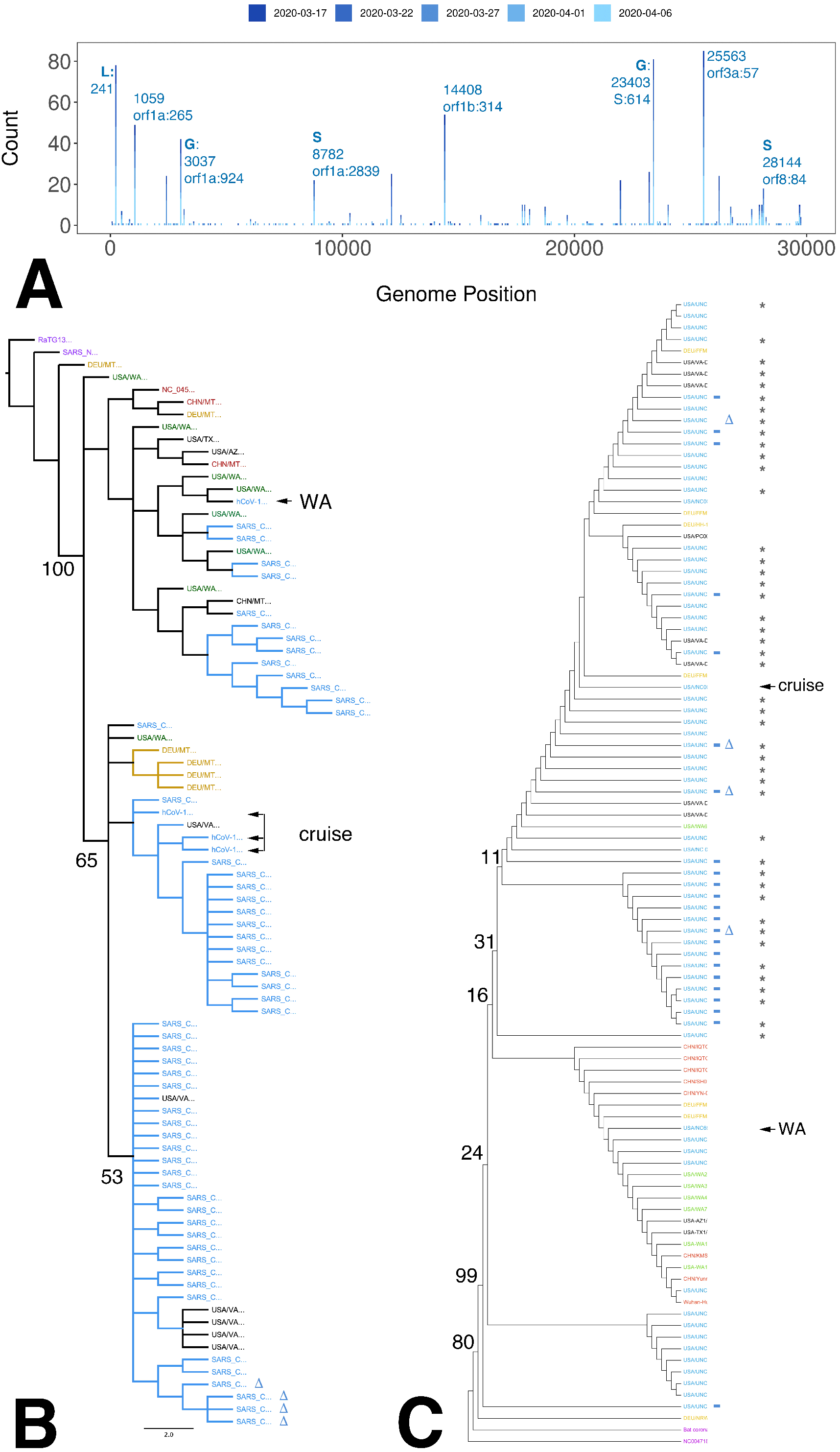
Phylogenetic analysis. A. Distribution of high confidence SNV across the genome. The genome positions (NC_045512) are on the horizontal axis and the count of samples on the vertical axis. Clade-defining SNV are indicated by GISAID designations. B. Neighbor-joining tree of whole SARS-CoV-2 genomes, including the first cases reported in NC: a person returning from Washington (WA) and a person participating in a cruise (cruise). The bat coronavirus genome strain RaTG13 was used as an outlier to root the tree. Average nucleotide difference is shown for the two major branches and the difference between SARS and SARS-CoV-2 (0.02). Colors indicate the approximate dates of disease. C. Neighbor-joining tree based on amino acids for S protein. Support values are listed at the major branch points. The colors indicate NC samples in blue, Washington State samples, including several independent sequences for SARS-CoV-2/Human/USA/WA1/2020, in green, representative other U.S. isolates in black, representative German isolates in gold, representative Chinese isolates, including NC_045512 in red. Additional genome sequences and protein sequences are from GISAID and GenBank.

Unlike retroviruses, such as human immunodeficiency virus (HIV), CoVs do next exist as sequence swarms, since CoV employ a proofreading RNA-dependent RNA polymerase (Agostini et al., 2018; Graham et al., 2012). Consistent with the biology of CoV, this study did not find widespread evidence of minor SNV.

Independently derived consensus genomes from the SARS-CoV-2/human/USA-WA1/2020 isolates showed evidence of divergence between the original isolate, the seed stock, and commercially distributed standard (**Figure 2B**). Similar culture-associated changes were recently reported for a second, culture-amplified reference isolate: Hong Kong/VM20001061/2020. This is not surprising, given that any large-scale virus amplification in culture is accompanied by virus evolution, but it raises concerns about the utility of using a natural isolate, rather than a molecular clone (Graham et al., 2018; Thao et al., 2020) as standard for sequencing.

The phylogeny based on whole genome nucleotide sequences revealed several interesting facets. Predictably, all UNC isolates of SARS-CoV-2 were significantly different from SARS-CoV and RatTG13 (**Figure 2B**, purple color). RatTg13 was used as an outgroup for clustering. The first NC case (NC_6999, (**Figure 2B**, arrow labeled “WA”)) was a person returning from Washington (WA) and sequence confirmed at the CDC (NC-CDC-6999). It initiated a branch of cases related to the original Wuhan isolate. The branch of cases (**Figure 2B**, arrow labeled “cruise”) contains the majority of NC cases, several cases isolated in neighboring Virginia (VA) (**Figure 2B**, black cases), and a cluster of cases reported in Germany (DEU, orange). It also contains several early cases, representing the individual that participated in a cruise.

SARS-CoV entry is determined by the spike protein ORF S and S is the target of neutralizing antibodies. **Figure 2C** shows the phylogenetic analysis of the S protein across all samples, the index cases for NC deposited by the NC Department of Health and Human services, and representative examples from the U.S., China, and Germany. Two branches emerged one containing isolates from China, Washington, and Germany, and a second containing U.S. and German sequences only. Since the S protein is shorter and more conserved across SARS-CoV-2, the limited numbers of SNV did not support as detailed a lineage mapping as the whole genome nucleotide sequences.

One large deletion was identified in four independent samples: 14 nucleotides were deleted beginning at position 29745 (indicated in **Figure 2C** by a delta symbol). This region is within the previously recognized “Coronavirus 3’ stem-loop II-like motif (s2m)”. This was confirmed in multiple isolates, supported by multiple, independent junction-spanning reads (**Figure 3A, B**). Junctions were mapped to single nucleotide resolution directly from individual reads. The variant 3’ end does not destroy overall folding but introduces a shorter stable hairpin (**Figure 3C, D**). How this mutation affects viral fitness remains to be established.

**Figure 3:**
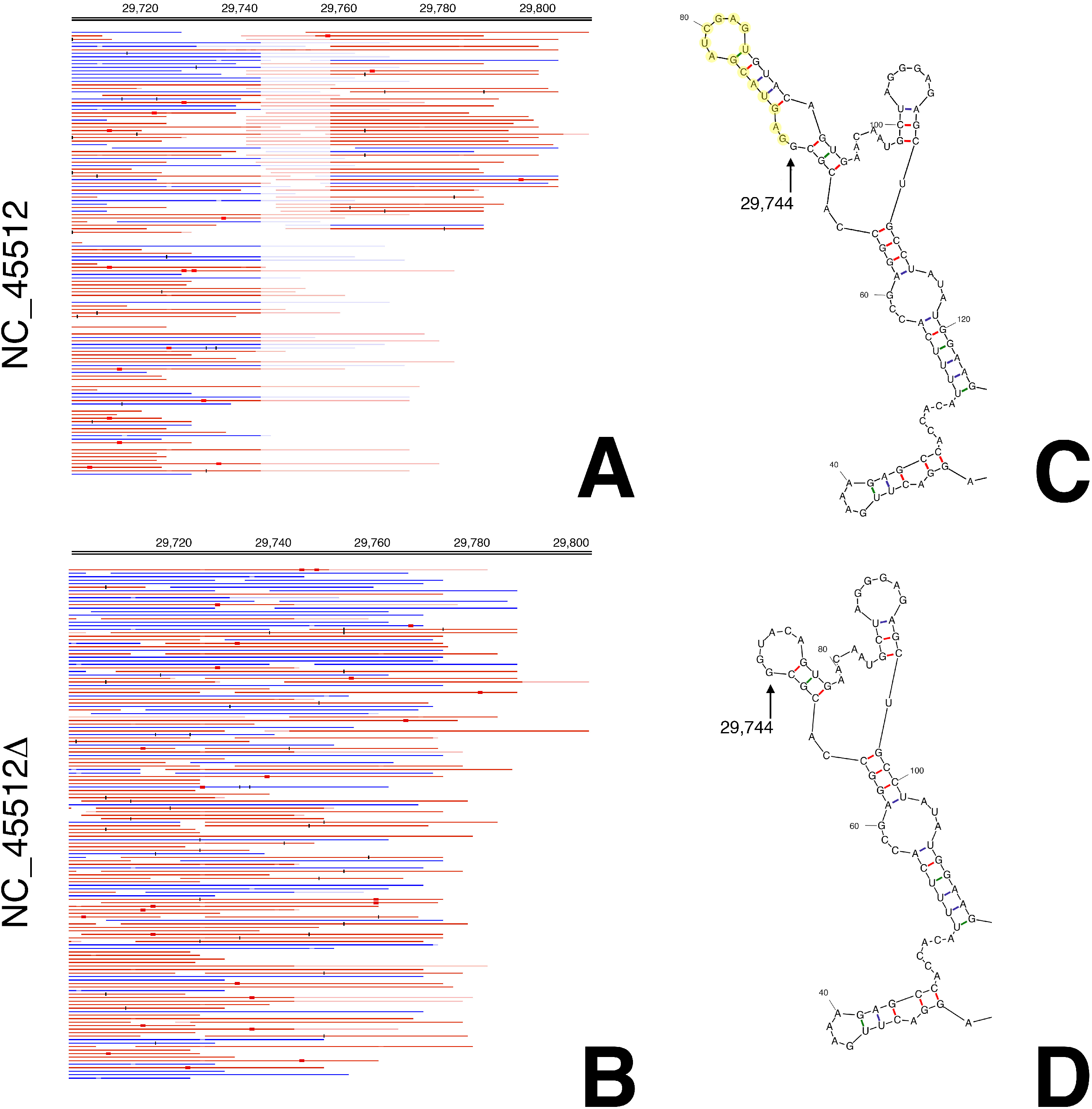
Detailed mapping of the variant 29745delta14. (A) Reads mapped to the reference sequence NC_045512. (B) The same reads mapped to an artificial target sequence with the 29745delta14. Blue indicates forward, red reverse reads (all reads are single reads). Red boxes and black bars indicated mismatches at below 20% reads (red) or above 20% of reads (black). In this alignment duplicate mapping reads were removed to guard against PCR amplification bias. Genome positions are shown on top (not that after nt 29745 the genome positions are out of sync due to the deletion). Note that this region is within the Coronavirus 3’ stem-loop II-like motif (s2m), annotated in NC_045512 as a prediction based on profile:Rfam-release-14.1:RF00164,Infernal:1.1.2. (C) predominant Mfold prediction of the 3’ end of NC_045512 with deletion bases indicated in yellow. (D) predominant Mfold prediction of the 3’ end of NC_045512 delta14.

In sum, this study generated exhaustive SNV information representing the introduction and spread of SARS-CoV-2 across a suburban low-density area in the Southern U.S. All samples were from symptomatic cases and the majority of genomes clustered with variants that predominate the outbreak in the U.S., rather than Europe or China. This supports the notion that the majority of U.S. cases were generated by domestic transmission.

## Discussion

This study demonstrates extensive shedding of SARS-CoV-2 in symptomatic patients among a low-density population in the southeastern U.S. It is the largest sequencing study that focuses on a suburban and rural community, rather than a crowded city, like New York City. The SNV distribution were consistent with continuous evolution or genetic drift of this new virus through an immunologically naïve host population (Consortium, 2004; Fauver et al., 2020; Lu, J. et al., 2020).

The first reported SARS-CoV-2 case in NC was a person who previously traveled to the state of Washington (03-03-2020, NC State Health Department). Additional early cases included person(s) who became infected while onboard a cruise ship (03-12-2020, NC State Health Department). Each of these introduction events was associated with a distinct clade. More recent cases, and cases in neighboring Virginia, were associated with the cruise case. This data supports the hypothesis that the majority of cases in NC originate from persons traveling within the U.S. rather than internationally, reflecting predominant spread by community transmission within the U.S. (Fauver et al., 2020).

SNV analysis documents the presence of a presumed high-pathogenicity variant D614G in 57% of the cases (Becerra-Flores and Cardozo, 2020; Ceraolo and Giorgi, 2020; Eaaswarkhanth et al., 2020). Within the limitations presented by measuring viral loads within samples collected at unknown times past infection and with presumably differing clinical sampling efficiency, patients with the D614G SNV presented with higher SARS-CoV-2 genome loads. Whereas the association of the D614G SNV with specific clinical presentations and high peak titers remains the subject of debate (Zhang et al., 2020)(doi.org/10.1101/2020.04.29.069054), it is clear that this variant signifies spread within Europe and the continental U.S. Given the increasing abundance of D614G SNVs, further research into its role in pathogenicity and clinical outcomes is warranted.

Four samples had the same 14 nt deletion in the 3’ UTR, and no samples had deletions within the coding region. This deletion is 71 nt away from the stop codon of orf10 (N protein) and eliminates a predicted stem-loop structure. An analogous bulged stem-loop at about the same location, right after the stop codon, is important for the replication of mouse hepatitis virus. In bovine coronaviruses an analogous RNA structure attenuates viral replication (Williams et al., 1999; Zust et al., 2008).

There seems to be partial overlap between the bulged stem-loop and the pseudoknot, suggesting that these two structures are mutually exclusive and may serve as a switch to regulate the ratio of full length RNA and defective RNA (Goebel et al., 2004). These two structures are also present in SARS-CoV. These isolates represent full-length genomes from symptomatic patients rather than disjointed RNA fragments recovered after clinical disease had subsided, thus we speculate that these deletion mutants are replication competent yet have altered ratio of full-length genomic and defective interfering RNAs. The biological phenotypes of these and other recent SNVs remain to be established through future studies.

There are limitations to our approach. These are similar to other NGS-based phylogeny reconstructions. Sampling was neither randomized nor exhaustive. At this point, we cannot exclude the presence of a founder effect and a disproportional impact of particular populations and situations on this dataset. The unknown group of samples were not “asymptomatic” in a broader sense of being negative for any respiratory symptoms. In the current time of limited personal protective equipment, limited sample kits, and limited testing capacity, it would not have been ethical to divert these resources for random population-wide sequencing. As properly randomized cohort studies become available in the future, the SARS-CoV-2 phylogeny will become more representative of SARS biology and less influenced by sample bias.

Some SNV may be the result of technical bias. For instance, the 5’ end awaits individual confirmation by RACE; the 3’ end likewise requires RACE for genome finishing. The Nextstrain database (Hadfield et al., 2018) suggests that positions 18529, 29849, 29851, and 29853 may be subject to PCR or sequencing bias. Lastly, targeted sequencing relies on amplification or hybridization capture. Unless the amplicon PCR primers or capture baits are completely removed a portion of reads will reflect the sequence that these primers/ baits were derived from rather than the sample. Most protocols rely on bioinformatic primer pruning alone. AmpliSeq, in addition to bioinformatic removal, enzymatically digests the targeting primers before library construction. Therefore, the sequences and SNV reported here could exclusively be attributed to the particular clinical sequence.

This study confirmed the sensitivity of current NATs with regard to the specific SARS-CoV-2 strains circulating in the region (and the U.S.). None of the UNC isolates had mutations in the CDC primer binding sites (Lu, X. et al., 2020). Three European isolates (MT358642, MT358639, MT318827) had a GGG>AAC polymorphism in the 5’terminal end of the forward CDC N3 (5’-<u>GGG</u>GAACTTCTCCTGCTAGAAT), which is a coronavirus consensus primer. Another European isolate (MT35638) had a G>T at 12,725 which is within the nCoV_IP2 forward primer. One European and one Chinese isolate (MT358638, MT226610) each had a SNV in nCoV_IP2 reverse primer at positions 12,818 and 12,814. As more and more viral genome sequences are generated, more and more SNV will be recorded including SNV in qPCR primer and probe binding sites. Currently, (May 9, 2020), 2.7% and 0.68% of sequences in GISAID contain SNV in the CDC primer pair N1 and N2, respectively. These data should be interpreted with caution, since at this point little standardization exists as to the quality of SNV reported and it is unclear how much a given SNV in one of the primer binding sites affects assay performance. Not all mutations in a primer binding site result in catastrophic failure or significant loss of sensitivity (Hilscher et al., 2005), which is defined as the sum of all steps in the assay pipeline, including, e.g. proper sample collection of the patient. Periodic re-testing of positive and negative samples by whole genome NGS represents an option to increase sensitivity and specificity and to detect any variants emerging in the populations, which may escape detection by NAT.

Testing by sequencing represents an interesting alternative to NAT in the case of coronaviruses, which are present at very high genome copy numbers during days of active shedding (Wolfel et al., 2020; Yu et al., 2020). Testing by SARS-CoV-2 targeted sequencing had perfect specificity, but lower sensitivity than qPCR (Sellers et al., 2020). Sequence coverage correlated with viral load. The lower sensitivity was expected as real-time qPCR amplicons can be placed anywhere on the target genome and optimized for sensitivity (Corman et al., 2020); shorter amplicons (< 100 base pair) maximize sensitivity as compared to larger amplicons (>200 base pairs) (Hilscher et al., 2005; Lock et al., 2010). By contrast, NGS represents a compromise as the entire viral genome has to be covered with primers that are part of a common pool. Primer design is governed by compatibility under a single set of conditions (annealing temperature) as much as by individual efficiency. The Arctic network protocol uses n = 96 larger amplicons (https://artic.network/ncov-2019). By comparison, the AmpliSeq protocol, deployed here, uses n = 237 amplicons of size 204±29 (mean ± sd), i.e. twice as many and substantially shorter amplicons with expected higher sensitivity. In sum, testing by sequencing represents a suitable, albeit expensive, tool for COVID-19 diagnosis.

About half of the specimen not clinically tested for SARS-CoV-2 tested positive by sequencing. This was not surprising, as to this day testing capabilities are limited, and probable cases are triaged based on clinical and public health indications. These unknown cases were not asymptomatic but represent patients with a clinically indicated need for upper respiratory sampling. Finding additional SARS-CoV-2 cases in this population suggests that case counts based on NAT represent a lower estimate of SARS-CoV-2 prevalence. It may also suggest that the current triage criteria for SARS-CoV-2 testing are too limited to understand spread of this virus. In sum, this study underscores the sensitivity and accuracy of current NAT assays and demonstrates the utility of testing by sequencing. It contributes to the worldwide effort to understand and combat the COVID-19 pandemic by providing the first set of full-length SARS-CoV-2 genomes from a non-urban setting.

## Acknowledgements

This work was funded by public health service grants CA016086, CA019014, and CA239583 to DPD. Funding was also provided by the University Cancer Research Fund and the UNC School of Medicine. The authors would like to thank all the members of the Damania and Dittmer labs, Corbin Jones and Nicole Fischer for critical reading, comments and suggestions.

## Figure Legends

**Supplemental Figure S1:**
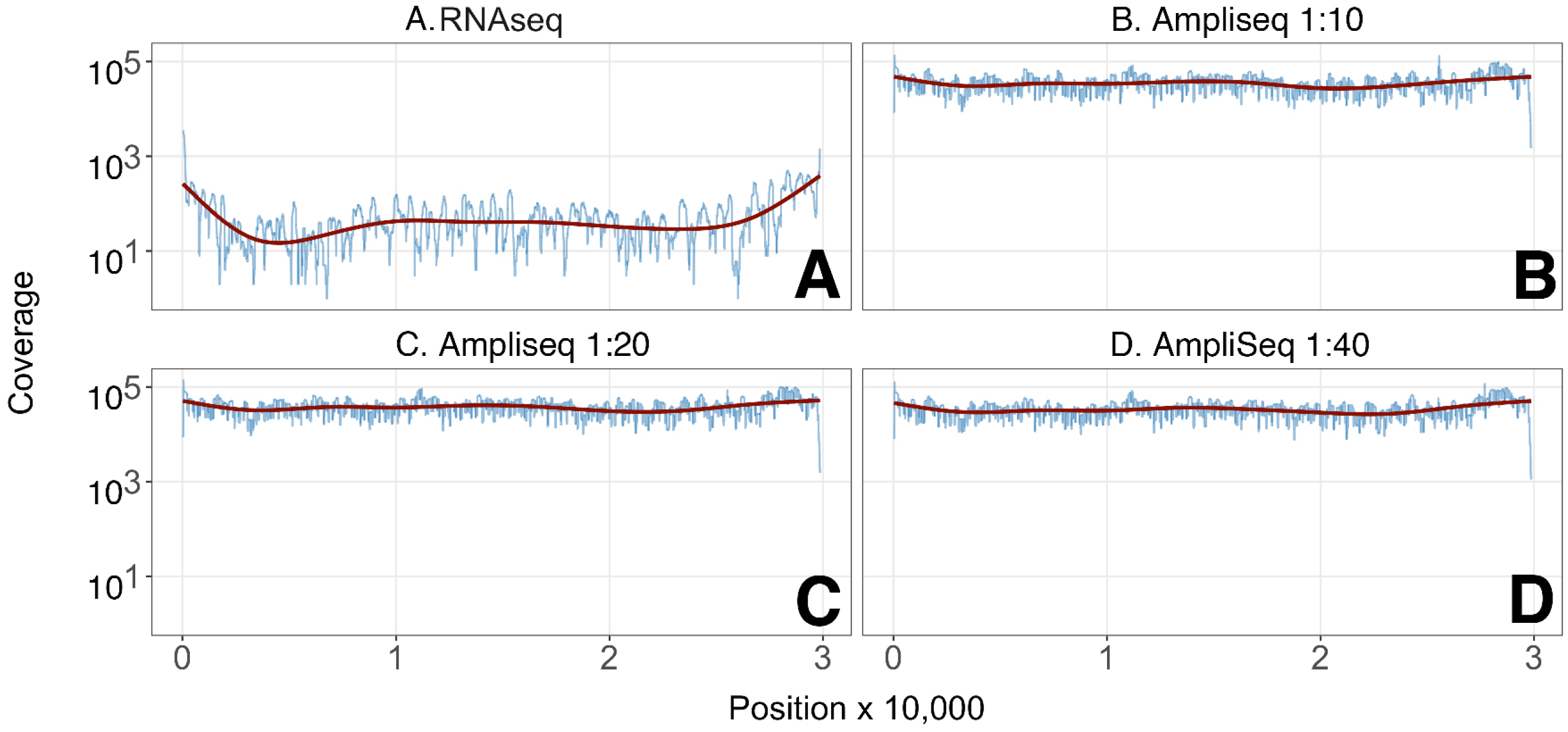
A-D. Coverage comparison of targeted (AmpliSeq) and non-targeted sequencing of the BEI reference material (NR-52285, strain SARS-CoV-2/human/USA-WA1/2020). Sample types, RNA seq or dilutions of input RNA, are listed on top. Coverage is shown on the vertical and genome position on the horizontal axis; Loess-regression line is shown in red.

**Supplemental Figure S2:**
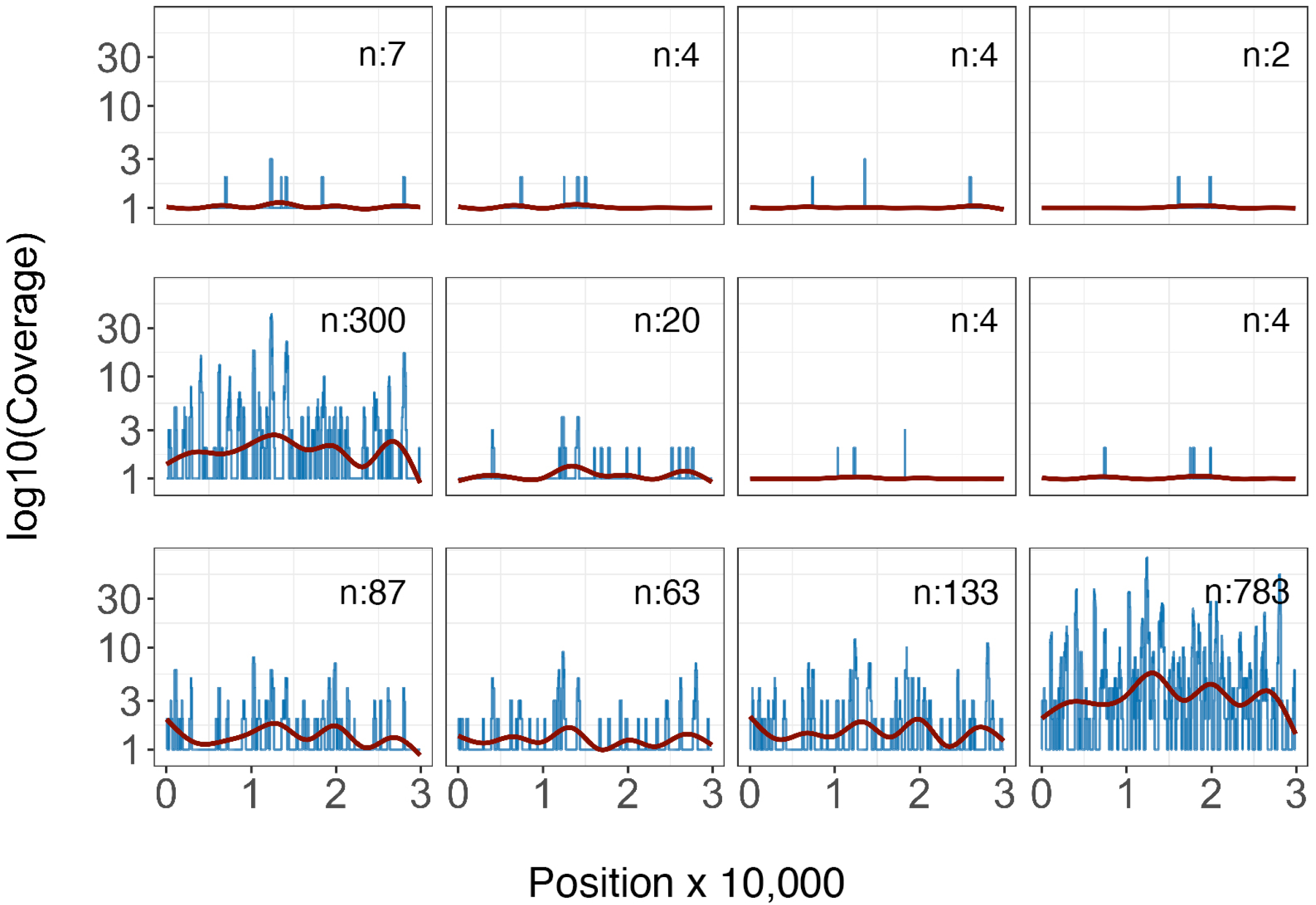
Coverage analysis of unknown cases which had at least one read mapped to the reference sequence NC_045512. The number of reads is shown on the vertical axis and genome position on the horizontal axis; Loess-regression line is shown in red. The insert label indicates the total number of reads.

**Supplemental Figure S3:**
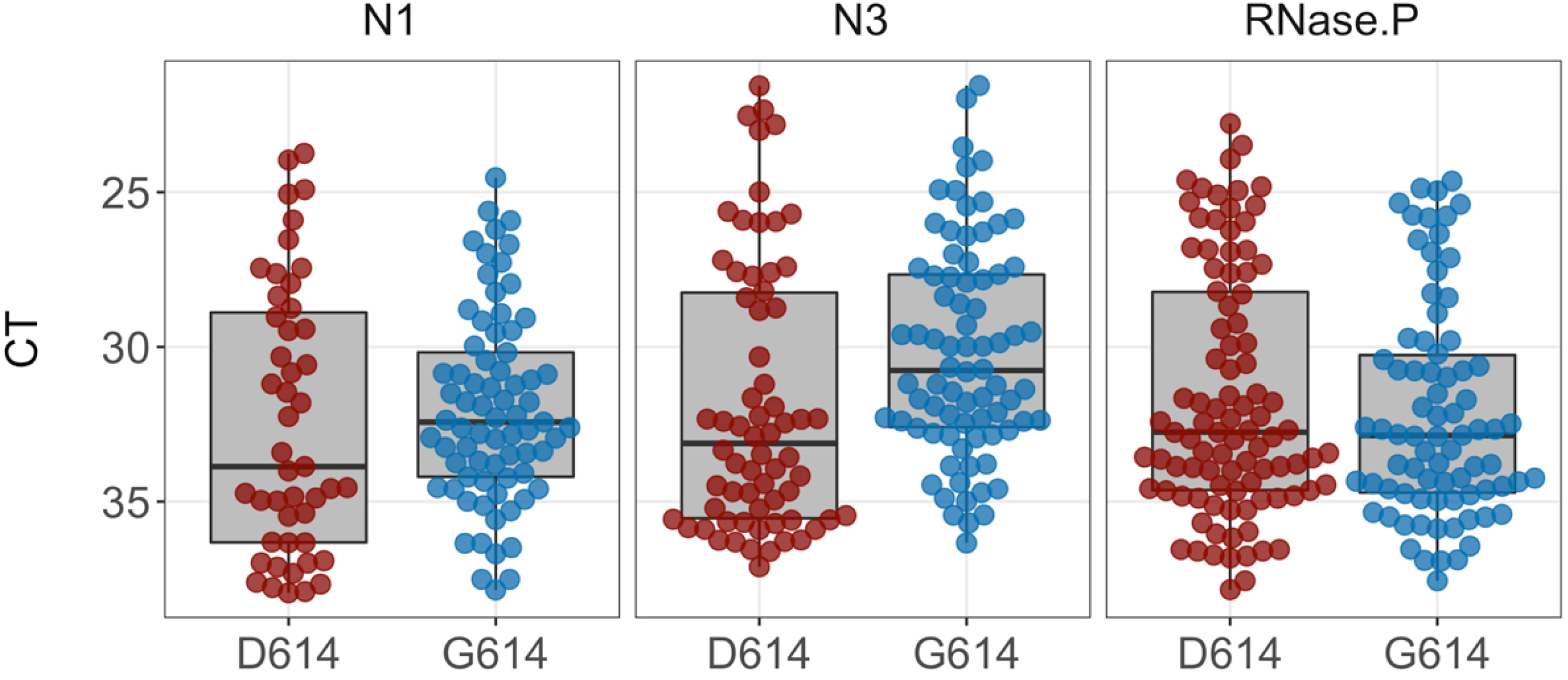
Beeplot of Raw CT numbers obtained by real-time RT-qPCR using CDC primers N1 and N3, as well as RNAseP, which serves as control for reverse transcription. Lower CT values signify higher genome copy number per sample. CT values are shown on the vertical and the SNV variant G614D (ancestral, red) or G614G (recent, blue) on the horizontal axis.

## Notes

### Competing Interest Statement

The authors have declared no competing interest.

